# A single-cell strategy for the identification of intronic variants related to mis-splicing in pancreatic cancer

**DOI:** 10.1101/2023.05.08.539836

**Authors:** Emre Taylan Duman, Maren Sitte, Karly Conrads, Adi Makay, Fabian Ludewig, Philipp Ströbel, Volker Ellenrieder, Elisabeth Hessman, Argyris Papantonis, Gabriela Salinas

## Abstract

Most clinical diagnostic and genomic research setups focus almost exclusively on coding regions and essential splice sites, thereby overlooking other non-coding variants. As a result, intronic variants that can promote mis-splicing events across a range of diseases, including cancer, are yet to be systematically investigated. Such investigations would require both genomic and transcriptomic data, but there currently exist very few datasets that satisfy these requirements. We address this by developing a single-nucleus full-length RNA-sequencing approach that allows for the detection of potentially pathogenic intronic variants. We exemplify the potency of our approach by applying pancreatic cancer tumor and tumor-derived specimens and linking intronic variants to splicing dysregulation. We specifically find that prominent intron retention and pseudo-exon activation events are shared by the tumors and affect genes encoding key transcriptional regulators. Our work paves the way for the assessment and exploitation of intronic mutations as powerful prognostic markers and potential therapeutic targets in cancer.

**Graphical Abstract:** 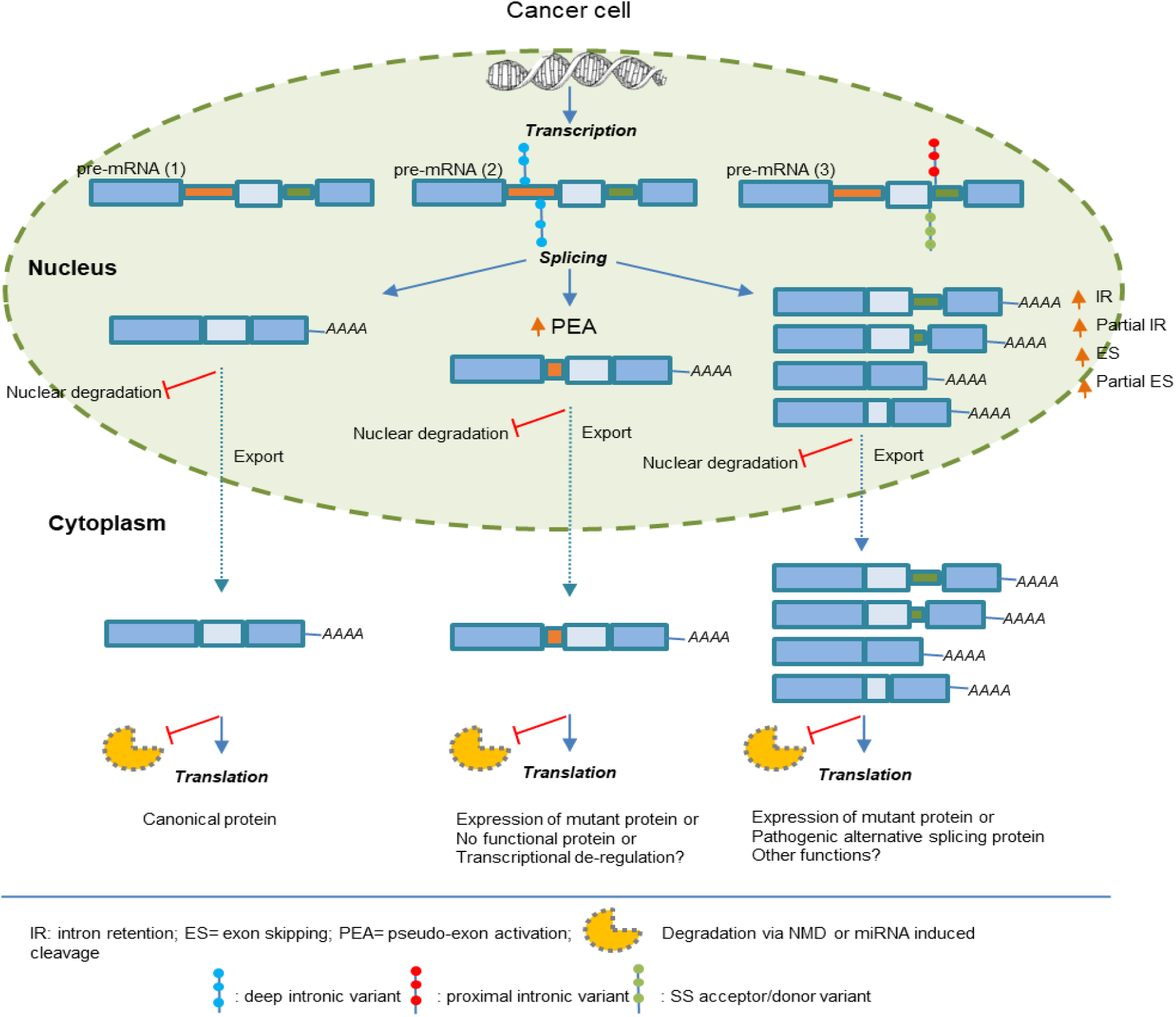

## Introduction

Introns in human genes can be extraordinarily large (up to 1 Mbp, with ∼3,400 being longer than 50 kbp and ∼1,200 longer than 100 kbp), and account for half of the non-coding human genome. As mutations in introns do not directly affect protein-coding sequences, they are usually overlooked ^1–3^. As a result, little attention is paid to the importance of intronic-located splicing regulatory elements that control the fidelity of pre-mRNA splicing and transcription timing. This is surprising given that pathogenic variants cause abnormal splicing changes, typically by damaging existing splicing motifs or creating novel splicing motifs and may comprise 15–60% of all human disease variants ^1–3^.

Recent studies have reported significant numbers of intronic variants and deletions in protein-coding genes that are associated with under- or overexpression of the affected genes or of distant genes interacting in 3D, thus influencing their regulation in normal or pathogenic conditions^4–6^. It is also well established that introns contribute to the control of gene expression by including regulatory regions and non-coding (yet functional) genes or even directly by their extensive length^6^. A substantial number of pathogenic variants located deep within introns (i.e., >100 bp from an exon-intron boundary) were recently reported, suggesting that sequence analysis of full introns may help to identify causal mutations for many undiagnosed clinical cases^4–9^. Moreover, direct associations between intronic mutations and certain diseases have also been reported, albeit sporadically^7–9^. These results agree with the findings we present here. For instance, we identified genes that are simultaneously overexpressed in basal-like cells from pancreatic cancer (PDAC) tumors and rank as the most mutated transcripts, particularly when we consider intronic variants.

However, there exist many limitations when investigating intronic mis-splicing variants, the main ones being the lack of approaches capable of simultaneously interrogating genomic and transcriptional information, and the lack of guidelines designed for assessing intronic variants and their contribution to abnormal splicing changes in disease. We address these limitations by introducing a novel pipeline that utilizes full-length single-nuclei and bulk RNA sequencing strategy for the “deep” characterization of genetic variability within introns and of their effects on splicing and gene expression in PDAC. Indeed, cancer is one of those diseases, where alternative splicing is the basis for the identification of novel diagnostics, and therapeutic strategies for therapy (e.g., antisense oligonucleotides or small-molecule modulators of spliceosome^9–11^). This new approach therefore holds promise for both the elucidation of fundamental biological principles connected to splicing regulation, and the identification of therapeutic targets in human disease.

## Results

### Strategy for the identification of intronic variants affecting splicing in PDAC

One of the most critical post-transcriptional mechanisms reprogramming transcriptional output and proteomic diversity in cancer cells is the loss of splicing precision when removing introns from pre-mRNAs^12^. Consequently, many mis-spliced variants are instead targeted for nuclear degradation or for nonsense-mediated mRNA decay (NMD) and thus, only few annotated alternative isoforms correspond to the precursors of the proteins mapped by large-scale proteomics studies^13^

To investigate this mechanism, we applied a new pipeline using a full-length snRNA-seq approach to three primary PDAC tumors (TM16, TM27 and TM56) and matched tumor-derived preclinical PDAC models (i.e., organoids and subcutaneous patient-derived Xenograft, PDXs). We then sought to determine pathogenic intronic variants causing abnormal splicing in these patient samples (**Figure 1A**). All samples were part of a unique patient cohort recruited for the Clinical Research Unit 5002 (https://gccc.umg.eu/en/cru-5002/).

**Figure 1:**
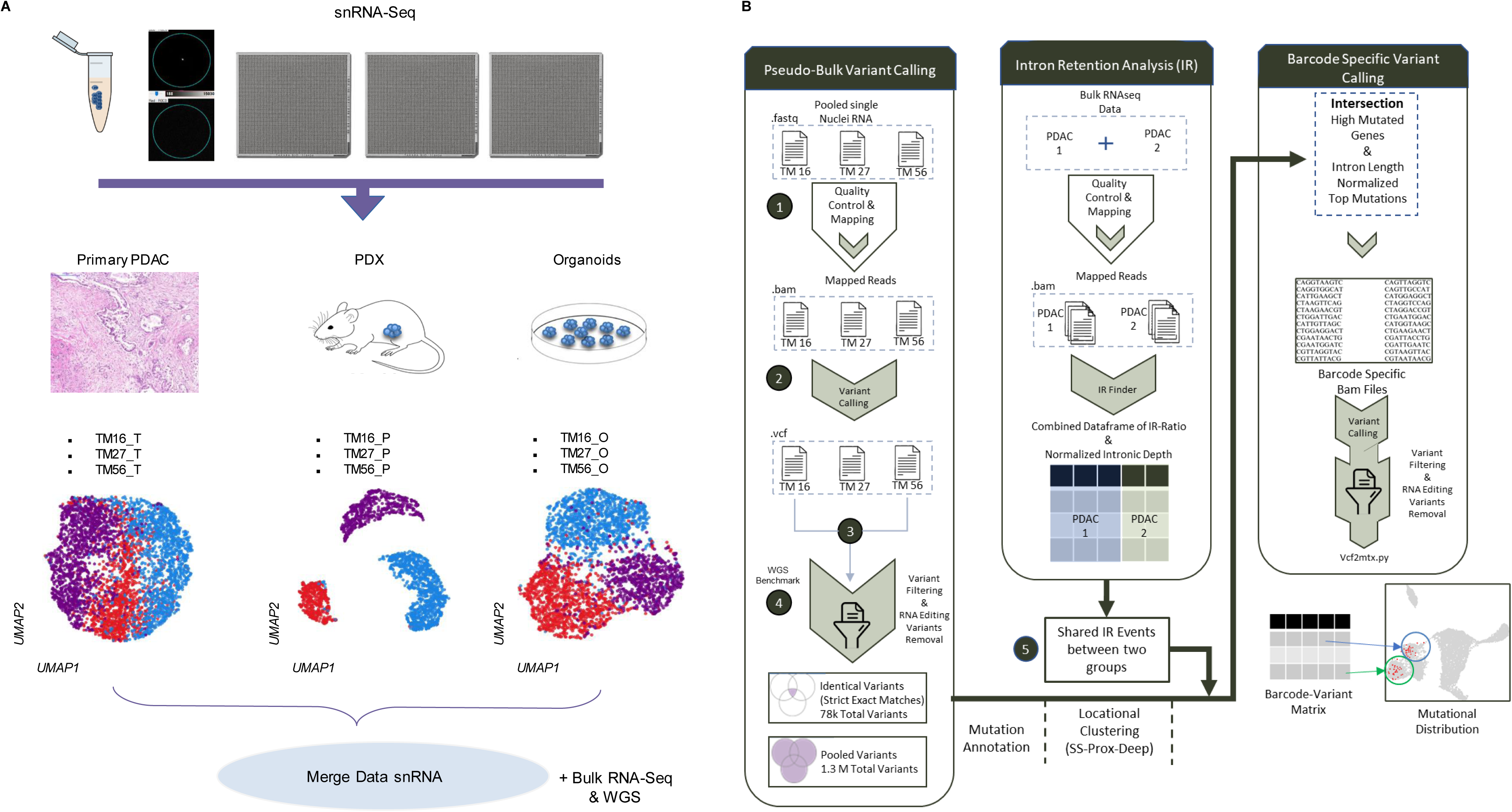
Strategy for the identification of intronic variants related to splicing in PDAC. **A**, snRNA-seq using the ICELL8 platform and the SMART full-length chemistry allowing for full transcript coverage was applied to PDAC primary tumors (TM16, TM27 and TM56) and matched organoids and PDX lines. A total of nine 5184-nanowell chips were performed (three for primary PDAC, three for organoids and three for PDX respectively and visualized in the UMAP. Finally, data from all nine nanowell chips were merged. Bulk RNA sequencing was performed additionally for each of the samples. **B**, DeepVarSplice variant calling pipeline combining snRNA-seq and bulk RNA-seq data determining intronic variant positions in the genome and intron retention events. 1) Gold standard RNA-seq variant calling pipeline pre-processing; 2) sample-level variant calling; 3) combination strategies for variant discovery (Shared-Combined); 4) Further filtering for RNA-editing variant removal; and 5) determination of shared intron retention events with weighted IR-ratio calculation between public and own PDAC samples.

For our snRNA-Seq experiments, we used the ICELL8 platform previously established in our group connecting genotype to phenotype in individual cells^15^. In contrast to the chemistry used by droplet-based platforms (i.e., in 3’-end approaches), ICELL8 is based on the SMART full-length chemistry allowing for the full read coverage of transcripts. Notably, when single nuclei are processed with this platform and chemistry, a strong enrichment in pre-mRNA is observed, including comprehensive coverage of introns and exons along these pre-mRNAs (**Figures 1A and 2C**). Furthermore, utilization of full-length chemistry allowed strong detection of intergenic and intronic sequences, as well as of non-coding RNAs, especially long intergenic non-coding RNAs (lincRNAs)^14, 15^.

**Figure 2:**
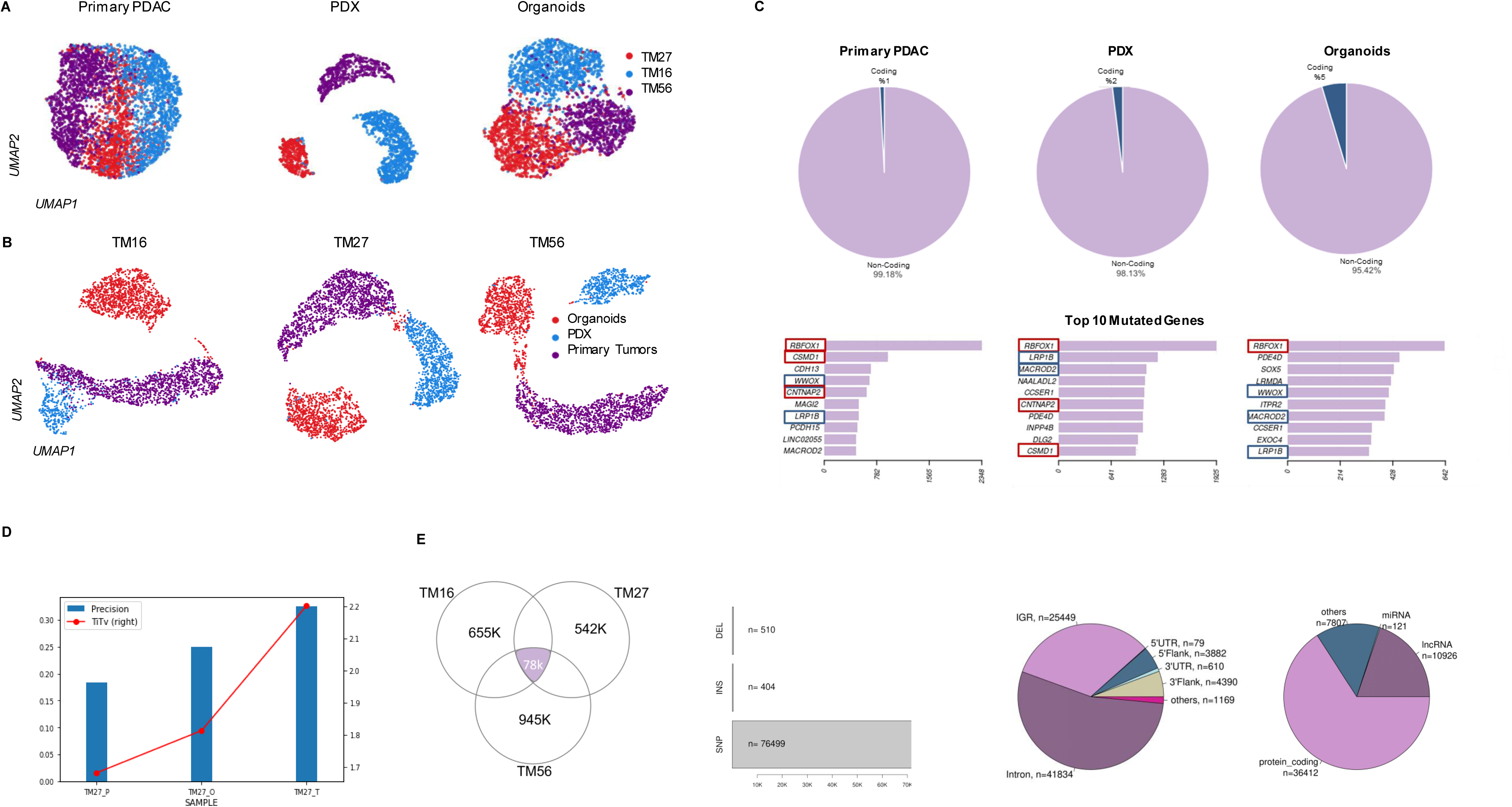
Identification of intronic variants using snRNA-seq data. **A**, UMAP plot for three primary PDAC tumors and corresponding models (TM16, TM27 and TM56). **B**, UMAP plot showing clustering by patients (primary PDAC and models) forTM16, TM27 and TM56. **C**, Pie charts of the three PDACs (tumor, organoids and PDX) showing percentage of exonic and intronic mutations (above) and bar charts represent number of top intronic mutated genes (below). **D**, Ti/Tv ratio of the variant analysis of primary PDAC and their matching models. **E**, Shared intronic variants and their classifications among all primary PDAC: TM16, TM27 and TM 56. Number of totals discovered and quality filtered intronic mutations from each sample (left), distribution of variant types among shared (78K) intronic mutations (middle), pie charts showing the distribution and location of the intronic variants in intronic, intergenic and coding regions, and a pie chart showing the intronic variants detected on coding and non-coding transcripts(right).

For the identification of intronic mis-splicing variants, we developed a variant-calling pipeline called “DeepVarSplice” that combines i) snRNA-seq data determining the variant’s position in the genome with ii) bulk RNA-seq data mainly capturing mRNAs and thus, charting splicing events in our samples (**Figure 1B**).

The pipeline begins with a pseudo-bulk snRNA-seq variant calling analysis using the per-sample GATK method (**Figures 1B (1-4)**), followed by an intersection of the individual findings to identify exact mutations present in all samples. The variants are then classified according to their genomic position considering intergenic, intronic and exonic regions using the maf file (Mutation Annotation File). By filtering the intronic variants, the genes were normalized and ranked according to intronic mutation load. The intronic variants are then forwarded to two parallel branches of the pipeline.

The first branch is set up for the investigation of these variants on the single cell level. Therefore, barcode-variant matrices containing the number of mutations for each gene (columns) and cell (rows) have been created to allow further transcriptional analysis related to the variants.

Simultaneously, the second branch of the pipeline connects genes that exhibit intronic mis-splicing to pathogenic splicing events through the investigation of partial or total intron retention (IR) or pseudo-exon activation (PEA) using the IRFinder algorithm (**Figure 1B (5)**). Finally, the intron mutated genes from the snRNA data were merged with the intron retained genes from bulk RNA-Seq data. As an outcome, we highlight genes showing mis-splicing related intronic variants that contribute to malignancy and thus proposed as potential therapeutic targets in pancreatic cancer.

To ensure the reliability of the variants detected by DeepVarSplice, we conducted comprehensive whole-genome sequencing (WGS) on two primary PDAC samples, namely TM27 and TM56. Our analyses encompassed gold-standard variant discovery, rigorous filtering, and benchmarking, utilizing both WGS and snRNA-Seq data, as depicted in **Supplemental Figure 1**. Out of the 153,000 shared variants detected by DeepVarSplice (snRNA-Seq data) in the two primary PDAC samples, 127,000 (83%) were confirmed using stringent criteria for variant calling in the WGS dataset. Both methods exhibited higher precision, with a Ti/Tv ratio of 2.3 for the snRNA-Seq data and 2.01 for the WGS data, thereby affirming the trustworthiness of the DeepVarSplice method.

### Identification of PDAC intronic variants using DeepVarSplice on snRNA-seq data

UMAP visualization of single nuclei clustering from the primary PDAC tumors, organoids and PDXs (**Figure 2A**) was strongly influenced by the patient of origin of each sample. To evaluate the consistency in transcriptomics and intronic mis-splicing variants, we merged data from all PDAC models of each patient and regenerated separate UMAP plots for TM16, TM27 and TM56 (**Figure 2B**). Then, by applying the DeepVarSplice pipeline (**Figure 1B**) to this snRNA-seq data, we observed that >95% of the mutations we identified were in non-coding sequences (i.e., in intronic or intergenic regions) and only ∼5% were found in protein-coding sequences. The most mutated genes at the level of introns included *RBFOX1*, *CSMD1*, *WWOX*, *CNTNAP2*, and *LRP1B* and were notably shared among PDAC tumors and models (**Figure 2C**).

To identify *bona fide* intronic variants, we determined the Ti/Tv ratio in each primary tumor-, organoid- or PDX-derived dataset. Interestingly, primary tumor data showed higher precision and a more realistic Ti/Tv ratio (2.18) in comparison to organoid- and PDX-derived data (**Figure 2D**). We therefore decided to focus on the three tumor-derived datasets for all ensuing analyses, which we complemented with bulk RNA-Seq data generated for 24 primary PDAC tumors from the CRU5002 cohort, as well as with public normal and tumor pancreas tissues data (accession ID: GSE211398).

In the end, variants shared between all three-tumor snRNA-seq datasets amounted to ∼78,000 non-coding variants with 41,834 in introns, 25,449 in intergenic regions (IGR), 4,390 in 3’ and 3,882 5’ gene flanks, 610 in 3’ UTRs, and 79 in 5’ UTRs. All shared variants detected in the primary PDAC with QUAL>30 is listed in **Supplemental Table 1**. Notably, 90% of these variants (69,488) were reported in the dbSNP_RS database based on WGS data remarking the validity of the variant calling performed with the snRNA Seq data. Moreover, the remaining 7,926 intronic variants identified as novel ones underscore the potential of snRNA-Seq datasets to detect both validated intronic variants and novel ones, demonstrating the added value of our approach.

Of these, 98.82% (n=76499) qualified as SNPs, 0.66% (n=510) as deletions, and 0.52% (n=404) as insertions (**Figure 2E**). Regarding the genomic annotation, most (n=36,412) mapped in protein-coding genes or lncRNAs (n=10,926) and very few (n=121) in miRNAs (**Figure 2C** and **Supplemental Table 1**).

### Classification and validation of intronic mis-splicing variants based on location

Next, to classify intronic variants we first performed a ranking based on the number of variants detected per gene normalized by the length and number of introns in each gene. Then, to link intronic variants to mis-splicing, we stratified mutations located 1–2 and 3–20 nt away from the nearest exon-intron junction, which we classified as donor/acceptor sites (SS) and branchpoint-proximal regions (BPs), polypyrimidine tracts (PPTs). All other variants >20 nt are categorized as “deep intronic”^5^. (see **Table 1** and **Supplemental Table 2**). Based on these indexes, many genes showed a high number of deep intronic variants, and some of these were reported as tumor suppressors with a multitude of alternative splicing variants expressed in specific tissues or only in tumor cells, e.g., *LRP1B*, *CSMD1*, *WWOX*, *FHIT*, *MTUS2, MAPK4 and MAP3K14* ^16–23^ (**Figure 3A**). The most prominent of these was *RBFOX1* and showed 208 deep intronic variants. The *RBFOX* family of RNA-binding proteins is well known to regulate alternative splicing (AS)^24, 25^. Recently, *RBFOX2* was reported to modulate a metastatic AS signature in pancreatic cancer^26^.

**Figure 3:**
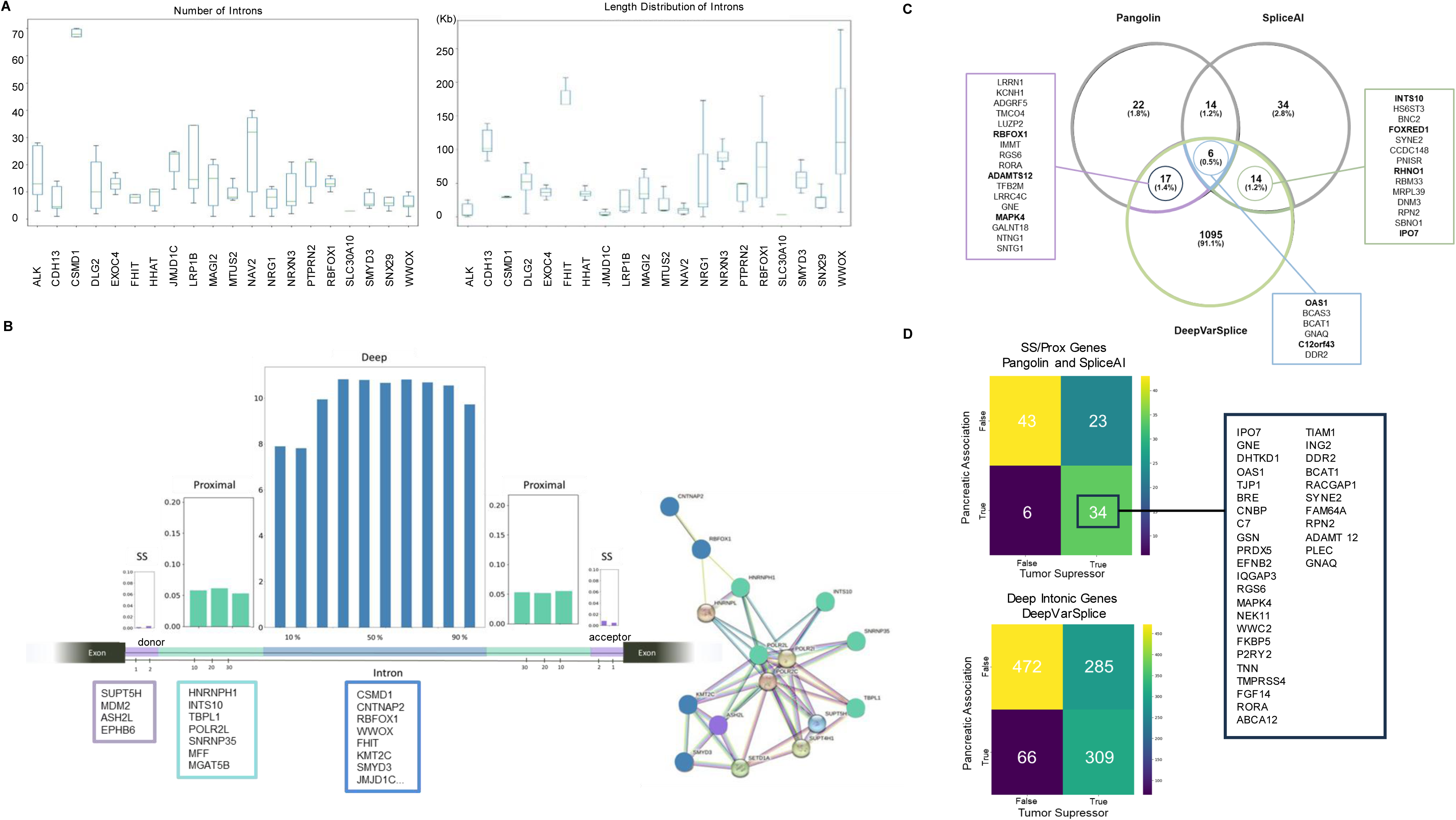
Selection and classification of genes exhibiting highest numbers of intronic variants. **A,** Boxplot showing the distribution of intron length and number of introns of top 20 genes with intronic mutations. **B**, Mutation percentages of all discovered mutations based on the variant locations (SS, Prox, Deep) and enriched pathway network representation with identified mutations. **C,** Performance of deep learning methods for mis-splicing detection: Comparison of SpliceAI and Pangolin and DeepVarSplice **D,** Pubmed Annotation of 107 Proximal and SS intronic variants scored by SpliceAI and Pangolin (top), 1132 Deep variants scored by DeepVarSplice (bottom).

**Table 1:** Classification of high intronic-mutated genes based on variant location. Classification of highly intronic mutated genes based on the variant location: donor and acceptor splice sites (SS, 1–2 bp away from the nearest exon-intron junction); proximal site (3–20 bp away from the nearest exon-intron junction), and deep intronic (>20 bp away from the nearest exon-intron junction). Total number of intronic mutations based on genes described in Supplemental Table 2.

We next examined the fraction of intronic variants over all variants at each position (**Figure 3B**). 99.6% of them were located deep in introns. Variants located in the proximal region are 0.32% of the total, with just 0.02% near SSs. By performing functional network analysis on the genes enriched for SS, proximal, and deep intronic regions, we identified enrichment of two main biological processes. The first, mRNA capping (GO:0006370), is most related to genes with variants in the SS and proximal region (e.g., *HNRNPH1, INTS10*, *TBPL1*, *POLR2L*, and *SNRNP35*). The second, regulation of transcription through histone methylation and H3K4-specific histone methyltransferase activity (GO:0051568), involved genes mainly carrying deep intronic variants (like *SMYD3*, *KMT2C*, *JMJD1C*, and *ASH2L*). Both GO terms are connected and the genes they contain are essential regulators gene expression and splicing ^27^. Finally, visualization of the distribution of intronic variants in top mutated genes *WWOX*, *SMYD3*, *JMJD1C*, and *NAV* (**Supplemental Fig. 2**) exemplifies read coverage from snRNA-seq and bulk RNA-seq in primary PDAC tumors and how these can be superimposed to evaluate splicing events.

Notably, DeepVarSplice identified 1,132 genes showing a substantial density of intronic variants that could potentially be linked to abnormal splicing. These genes were selected based on the modified Z-score method. This method is particularly apt for the normalized variant numbers dataset, which exhibits a non-normal distribution. The modified Z-score uses the median and the Median Absolute Deviation (MAD), providing robustness against skewed distributions (**Figure 3C**).

To validate our findings, we also employed recently developed Deep Learning-based tools, specifically i) Pangolin and ii) SpliceAI ^28–29^ designed for detecting potential intronic variants causing pathogenic splicing.

Out of the 78,000 variants originally identified (**Supplemental Table 1)**, 295 variants affecting 107 genes were scored as potentially pathogenic by at least one of these tools (**Figure 3C**). Through a direct comparison involving Pangolin, SpliceAI, and DeepVarSplice, we identified 34 genes depicted as tumor suppressors in pancreatic cancer in **Figure 3D**. One explanation for the heightened sensitivity of variant detection using our approach is its ability to access variants deep within introns from the intronic regions of pre-mRNAs present in sn-RNA data. In fact, a limitation of SpliceAI and Pangolin is their optimization primarily for variants located within 50 base pairs on the splice site defined as SS and proximal intronic regions, affecting canonical splicing. Furthermore, it’s important to acknowledge that both Pangolin and SpliceAI were developed predominantly using bulk-RNA sequencing methods, leading to potential gaps in the dataset due to the absence of intronic sequences. This limitation highlights the advantage of our approach in capturing a more comprehensive spectrum of intronic variants.

### Transcriptional regulation relates to mis-splicing in basal-like tumor cells

We performed transcriptional subtyping of single tumor nuclei into the more aggressive/drug-resistant “basal-like” (BL) or the better prognosis-associated “classical” subtype (CLA) using a ranking markers method described previously ^30^ on snRNA-seq data. The distributions of each subtype in each tumor showed BL cells highly represented in primary tumors, and partially in matched PDXs. In contrast, CL cells predominated in our organoid models (**Figure 4A**). Next, we performed differential expression analysis (DEG) between BL and CL tumor cells, where we identified a few hundred markers for both subtypes with absolute log_2_FC >0.5 and *P*_adj_ <0.05 (**Supplemental Table 3**) and visualized the most prominent ones in UMAP plots (**Figure 4B**).

**Figure 4:**
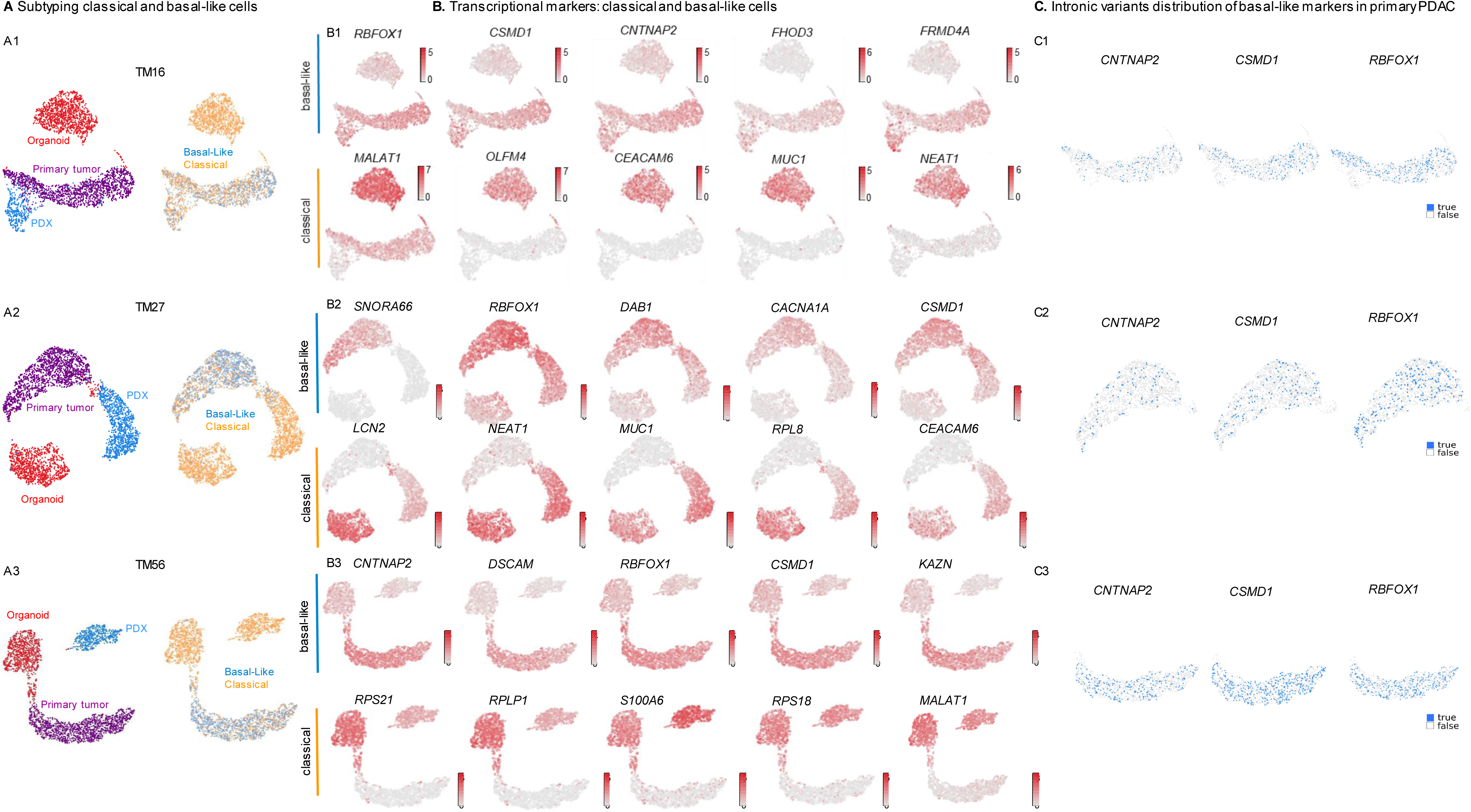
Tumor subtype and marker identification at the single-nucleus level. **A**, UMAP plot of single cells from the three patients colored by tumor model (left) and by inferred tumor subtype (right). **B**, UMAP plot showing the gene expression of the top transcriptional markers comparing classical vs basal-like cells. **C**, UMAP plot showing the mutated primary tumor cells for *CNTNAP2*, *CSMD1* and *RBFOX1*

Tumor heterogeneity was assessed by evaluating cell types using snRNA-seq from all models at hand. As expected, primary tumors exhibited the highest heterogeneity as regards cell type composition while organoids mostly contained ductal-like cells (**Figures 5A-C**). Strikingly, *RBFOX1*, *CSMD1*, and *CNTNAP2*, our topmost intronic mutated genes (**Figures 1B** and **4C**) appear to be simultaneously the genes most upregulated in BL cells (**Figure 5A**). At the same time, the markers found in CL cells have already been described as biomarkers for pancreatic cancer: *MALAT1*, *NEAT1*, *CEACAM6* or *MUC1* ^31–34^. For the lncRNA *NEAT1*, two novel mutations are reported here for the first time (**Supplemental Table 1**). Taken together, CL cells and models are characterized by less abundance and relevance of intronic variants according to the output of our pipeline, thus suggesting that mis-splicing mechanisms are linked to the aggressiveness of BL tumors (**Figure 5D**).

**Figure 5:**
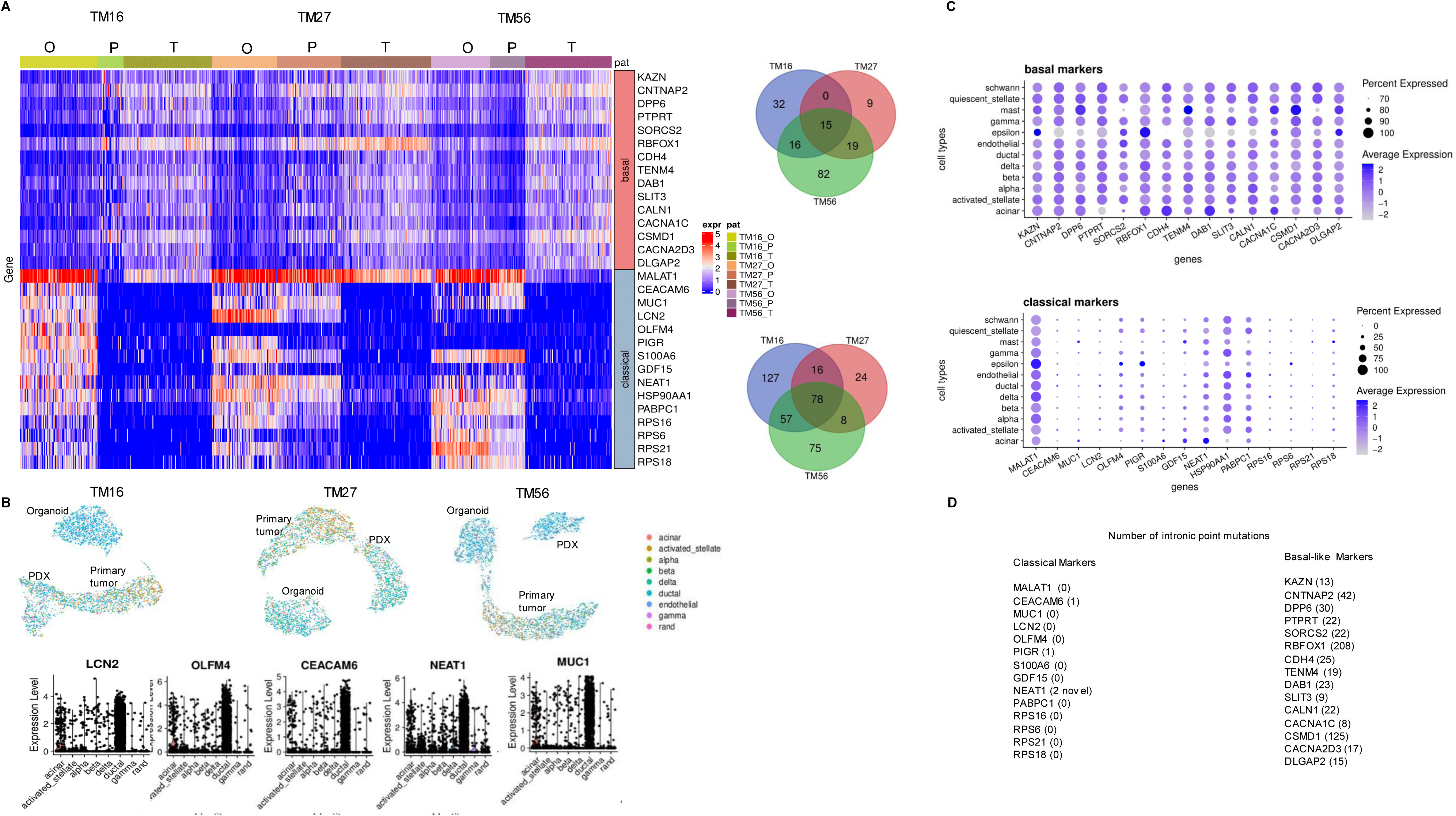
Identification of markers for BL and CL tumor cells. **A**, Heatmap showing gene expression of the top 15 basal-like and classical-like markers overexpressed in all PDAC. Venn diagrams illustrate the overlap of marker genes between tumor PDAC patient samples. **B**, Dot plot showing expression of the markers in each of the pancreas cell subtypes. Circle size is proportional to the percentage of cells in each cell type expressing the marker and circle color represents the average marker gene expression in the cell type. **C**, UMAP plots by patient showing clusters annotated to specific cell subtypes. The violin plots below represent five classical pancreas markers across cell subtypes. **D**, Table showing the number of intronic mutations of the top CL and BL marker genes.

### Contribution of intronic mis-splicing variants to pathogenic splicing and poor PDAC prognosis

In general, mis-splicing affects the regulation of genes (up- or downregulation) or generates new isoforms in normal and tumor tissues ^35–37^. However, our focus here is on the identification of potentially pathogenic splicing events in PDAC. It is important to note that the definition of pathogenicity in this study is based on comparisons to human references (via SNP calling) or to non-tumor pancreatic tissues when assessing mis-splicing findings. Nevertheless, we acknowledge that conclusive assertions regarding pathogenicity necessitate the inclusion of functional studies.

To determine intronic retention (IR), we employed the IRFinder tool by intersecting the findings of bulk RNA-seq from 24 primary PDACs recruited for CRU5002 and 9 public PDACs (GSE211398). To ensure robustness and significance, we included additional healthy and tumor pancreas tissues sourced from credible databases to visualize and verify the IRFinder results (SRA and GEO). A common challenge in utilizing public transcriptomic (bulk RNA-Seq) data for this research field is the limited sequencing depth, which often hinders precise IR event identification. To address this limitation proactively, we sequenced 24 primary PDAC samples from the CRU 5002 with higher depth, guaranteeing a minimum of 100 million reads per sample, specifically for splicing investigations resulting on a total of 2,489 shared IR events (listed in **Supplemental Table 4**).

Next, we take a closer look into those genes found previously to be i) most enriched for intronic variants (**Table 1**) and ii) simultaneously found overexpressed in BL tumor cells (*RBFOX1*, *CSMD1*, and *CNTNAP2* from **Figures 1B** and **4C**). All three are long genes with many (mostly small-, <100 nt, and micro-, <60 nt) exons, and with relatively small introns. Moreover, all intronic variants discovered within these genes were situated deep within the intronic regions, establishing an association with non-canonical splicing regulation. Among these genes we have made a noteworthy discovery of combinatorial abnormal splicing ^5^ showing multiple splicing events within a single gene. For example, we reported a combinatorial splicing in a small portion of the *CSMD1* gene (chr8: 3,187,000-3,202,000) showed an unusual exon skipping in the middle of the exon (chr8: 3,189,980-3,190,067) followed by two PEA events (chr8: 3,197,495-3,197,578 and 3,200,391-3,200,466; **Figure 6A**).

**Figure 6:**
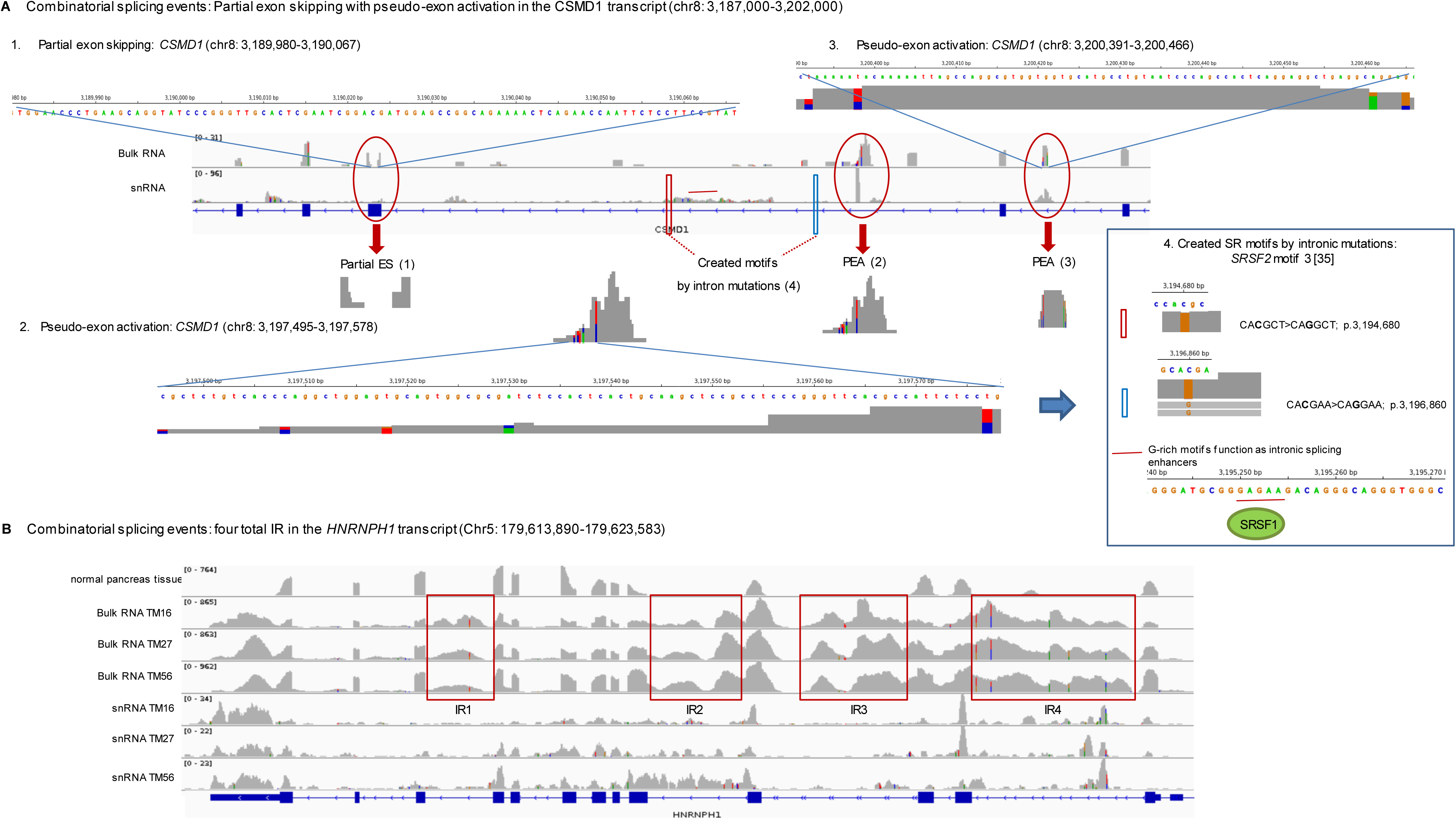
Combinatorial splicing events related to intronic variants in PDAC. **A,** Partial exon skipping with pseudo-exon activation in *CSMD1* (chr8: 3,187,000-3,202,000). 1: partial exon skipping in *CSMD1* (chr8: 3,189,980-3,190,067); 2: pseudo-exon activation in *CSMD1* (chr8: 3,197,495-3,197,578); 3: pseudo-exon activation in the *CSMD1* (chr8: 3,200,391-3,200,466). **B**, Four combinatorial IR events in *HNRNPH1* (Chr5: 179,613,890-179,623,583).

To assess potential non-canonical splicing mechanisms triggered by deep intronic variants in the *CSMD1* gene, we conducted a search for RNA binding motifs near the observed mis-splicing events. The presence of these variants generates motifs, such as SR-protein binding sites, leading to the activation of introns (PEA). Following motif analysis for SR-protein binding sites, we identified two intronic variants in our snRNA-seq data creating two *de novo* SRSF2 motifs^38^: i.e., C>G substitution: CA**C**GCT>CA**G**GCT; p.3,194,680 and CA**C**GAA>CA**G**GAA; p.3,196,860 (**Figure 6A**). This suggests that intronic variants are located deep in the intron and creating additional SR binding sites that may synergistically contribute to activate PEA events. To this day, few examples of PEA have been reported as caused by deep intronic mutations without directly changing a splice site sequence ^39, 40^.

In addition to genes with significant mutations deep within introns, our findings include other genes exhibiting high mutation rates in the SS and adjacent proximal intronic regions. One example is *HNRNPH1*, reported by our pipeline for carrying high number of intronic variants in proximal regions (**Table 1**). For HNRNPH1 we detected combinatorial splicing involving four IR events (**Figure 6B**). HNRNP nucleoproteins are known to associate with pre-mRNAs in the nucleus and influence their processing and other aspects of mRNA metabolism and transport ^41^.

Several transcripts encoding members of the Integrator complex (*INTS10*, *INTS3*, *INTS11*) were affected by multiple IR events (see *INTS3* examples in **Figures 7C and 7D**). As the Integrator complex interacts with the C-terminal domain of RNA polymerase II to allow processing of U1 and U2 small nuclear RNAs, these splicing alterations could indirectly affect splicing in PDAC. Closer inspection of the two IR events in *INTS3* revealed an enrichment for mutations in snRNA-seq data that give rise to CCC sequences that follow a GGG motif. These give rise to partially overlapping recognition motifs for SRSF1 and SRSF2 (**Figure 7E**) and could function as splicing enhancers to compensate for weak PPTs tracts^42^. The high intronic mutation load we uncovered using snRNA-seq suggests that the occurrence of such events at the single-cell level might be significantly more frequent than previously presumed_40._

**Figure 7:**
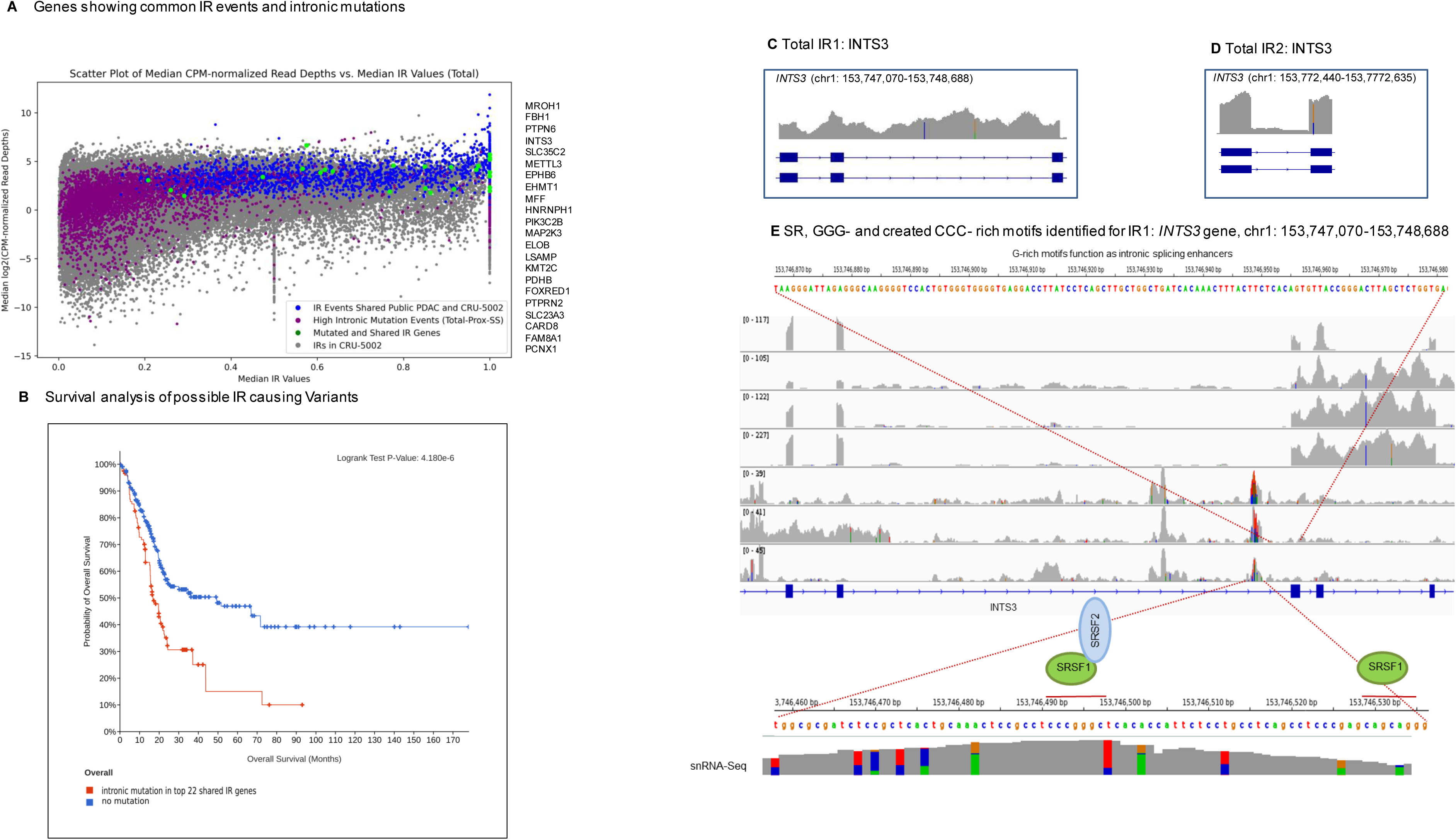
Analysis of IR related to intronic variants in PDAC. **A,** Scatter plot of IR ratios from CRU5002 bulk RNA-seq data with their corresponding normalized read depths. Filtered and shared IR events between public PDAC and CRU-5002 data (blue). IR events of high intronic mutated genes (Total-SS-Proximal; green). List of genes exhibiting IR events related to high intronic mutations (right). **B**, Kaplan-Meier survival analysis generated via the cBioPortal shows overall survival time of PDAC patients with or without mutations in the top 22 IR genes shared by all three tumors. **C**, Region of high percentage IR event detected by IRFinder for the *INTS3* gene shared in KFO and public PDAC (Mean IRraio: 0.45). **D**, Second IR event in the *INTS3* gene (Mean IRraio: 0.25). **E**, SR binding motifs identified in *INTS3*. Two CAGCAGG binding sites for *SRSF1* and one CTCCCGG motif for SRSF2.

Finally, we assessed the clinical outcomes and significance of intronic variants associated with mis-splicing in pancreatic cancer by combining the sn-RNA-seq from the PDACs with the bulk RNA-Seq data from the PDACs (sourced from the CRU5002 and public data). The scatter plot depicted in **Figure 7A** illustrates shared intronic retention (IR) events, represented in blue, between the bulk RNA-Seq data from the CRU5002 and public datasets. Additionally, genes with a high mutational rate in introns (SS-Proximal and deep), identified in the sn-RNA-seq data from primary PDACs using our pipeline, are shown in green. Consequently, from this analysis, we identify and highlight a subset of 22 candidates. Notably, survival analysis performed with the TCGA-PAAD cohort and generated by the cBioportal for the 22 genes revealed their significant association with poor prognosis in PDAC (**Figure 7B**). Six of these genes (*METTL3*, *HNRNPH1*, *INTS3*, *ELOB*, *EHMT1* and *KMT2C*) are linked to the GO terms of pre-mRNA capping (GO:0006370) and histone H3K4 modification (GO:0051568) previously described in **Fig. 3B** by performing the DeepVarSplice pipeline.

## Discussion

The current study proposes a new strategy for the investigation of intronic mis-splicing variants and their role in promoting pathogenic splicing in PDAC. In contrast to prior investigations utilizing whole-genome sequencing and transcriptomic data, our approach employs a comprehensive full-length snRNA-seq method, offering high-resolution identification of intronic variants. For this new method we developed a comprehensive pipeline, the DeepVarSplice that takes advantage of integrating multi-omics information e.g., variant calling, transcriptomics and splicing within a cell and thus offers a more holistic view of the underlying molecular mechanisms in complex diseases.

Simultaneously, we address the challenges associated with DeepVarSplice in handling low-covered regions, non-uniform read distribution and thus, increased false positive findings. To overcome these limitations, we employed gold standard methodologies, specifically conducting whole-genome sequencing to validate our SNP calling performances^43^ using sn-RNA-seq data. Additionally, we utilized well-established artificial intelligence tools (AI tools), including Pangolin and SpliceAI to affirm our results concerning intronic variants associated with potential pathogenic splicing. Notably, both Pangolin and SpliceAI exhibit limitations when assessing intronic variants deep within introns, as they were primarily optimized for variants in the splice site (SS) and proximal region (within 50 base pairs of the splice site). Additionally, these tools were originally developed using bulk RNA sequencing datasets. For the first time in this study, we employed both tools using our full-lenght sn-RNA-seq approach. Consequently, our DeepVarSplice pipeline demonstrated enhanced sensitivity in intronic variant detection, capitalizing on the presence of pre-mRNAs in sn-RNA data and achieving full coverage of both intronic and exonic regions within a gene. Lastly, it’s crucial to emphasize that, for both splicing analysis and variant detection using sn-RNA-seq approaches, a deep sequencing strategy is vital to enhance sensitivity and reliability of findings. Accordingly, we conducted 300 to 400K median reads per nucleus for snRNA-Seq and 100 to 200 million reads per sample, paired-end, with 300 cycles for bulk RNA sequencing.

Our approach delivers extensive information on the landscape of primary transcripts, especially near potential pathogenic splicing events. Thus, we can begin to decipher the complexity of RNA sequences acting as suppressors or activators of splicing in the context of PDAC. Recent high throughput characterization of exon splicing enhancer and silencer (ESE and ESS) motifs has indicated that pseudo-exons can be discriminated from genuine exons on the basis of their low ESE and high ESS contents ^44, 45^. Moreover, these motifs can be found both in exons and in introns, and function via the recruitment of sequence-specific RNA-binding proteins that can dictate splicing choices ^46–48^. Our analyses suggest that splicing dysregulation in PDAC can be linked to non-canonical sequence signals, i.e., to intronic variants affecting RNA motifs located deep in introns. For instance, we found several motifs for SR RNA-binding proteins (e.g., SRSF1 and SRSF2), enriched for GGG and CCC, distributed in the SS-proximal regions of introns and, in some cases, exhibiting extensive overlap (**Figure 7**) ^38, 48^. Although RNA-binding proteins like SR proteins bind to RNA with high sequence specificity, it is difficult to obtain well-defined consensus motifs for each of them ^38, 47–48^. However, cooperation and competition between the SRSF1 and SRSF2 proteins have been reported in the regulation of alternative splicing events, which are related to synergistic and compensatory interactions with target RNA ^35^. Still, the exact mechanism by which these variants affect the splicing machinery, and its downstream choices remains to be dissected, and very few examples of deep-intronic disease-causing mutations have been described to date ^39, 40^.

The complex nature of splicing events and their regulation necessitates the development and implementation of new approaches to uncover nuanced relationships between mutations and splicing choices. We described a pseudo-bulk variant-calling pipeline, the DeepVarSplice exploiting dense and full-length coverage of snRNA-seq datasets to detect putatively pathogenic deep intron variants and link them to mis-splicing events occurring in PDAC tumors, while also gauging cell heterogeneity. As our method generates a multidimensional output that typically serves as the foundation for machine learning models (ranging from Support Vector Machines to Convolutional Neural Networks), we envisage a near-future combination that would vastly improve our understanding of the combinatorial code regulating splicing choices in the context of cancer.

## Acknowledgements

We are grateful for excellent technical support provided by Jacqueline Fink and Waltraut Kopp, and to Hanibal Bohnenberger for technical advice and for providing patient-derived samples. We also thank Jeniffer Appelhans and Mark Bösherz for the generation of organoids derived from primary PDAC tumors.

## Author Contributions

G.S., A.P., and E.H. conceived the project and experiment design. V.E and P.S supported the experimental design and contributed to experimental data G.S. led the method development. F.L. led the experimental data production. A.M. contributed to experimental data, E.T.D. and M.S. led and performed the data analysis. K.C supported data analysis and A.P. and G.S. wrote the manuscript. All authors have read and agreed to the submitted version of the manuscript.

## Competing interests

The authors declare no competing interests.

### Institutional review board statement

The utilization and characterization of human PDAC data and samples within the CRU5002 have been approved by the ethical review board of the UMG (11/5/17).

### Informed consent statement

Informed consent was obtained from all subjects involved in the study.

## Methods

### Patients’ sample information

Utilization and characterization of human PDAC data and samples within the CRU5002 has been approved by the ethical review board of the UMG (11/5/17). Sequencing studies and the generation of organoids and PDX models have been performed using tumor tissue from CRU5002 PDAC patients with progressed disease upon histological PDAC confirmation.

### Generation of PDX models

For the generation of PDX models, bulk tumor tissue was subcutaneously transplanted into SHO-*prkdc^scid^Hr^hr^* mice. Engrafted subcutaneous tumors were passaged in mice for three generations prior to snRNA-sequencing.

### Generation of organoids models

Tumor tissue was minced and digested in Dulbecco’s modified Eagle’s medium containing 5 mg/ml of collagenase XI (Sigma-Aldrich, C9407), DNAse final concentration 10µg/ml (SIGMA D5025-150KU) and Y-27632 final concentration 10,5µM (Adooq Bioscience, 129830-38-2) and incubated at 37°C for 45 min. The material was further embedded in Matrigel (Corning, New York, USA; Cat#356231) and cultured in human pancreatic cancer complete medium (Wnt3a, R-spondin1, Gastrin, hEGF, A 83-01, hFGF-10, mNoggin, Primocin, N-acetylcystein, Nicotinamide, B27 supplement and Y-27632). For passage, the Matrigel-containing organoid was digested by TrypLE™ Express (Thermo Fisher, 12605-028) with DNAse and Y-27632 as described above for 15 min. The sample was centrifuged at 500g for 5 min, and the precipitated cells were embedded in GFR Matrigel and cultured in human pancreas organoid complete feeding medium.

### Nuclei extraction from primary PDAC and PDAC models

Nuclei isolation of PDAC was performed according to the “Nuclei Isolation from Cell Suspensions & Tissues for Single Cell RNA sequencing”, Document Number CG000124 Rev E, 10x Genomics, (2021, June 30). For organoids, minor modifications were performed. The cells were spun down at 500 x g, lysis took place for 15 minutes and only two washing steps were performed. Following the second wash, the nuclei pellet was resuspended in 1 ml wash buffer in preparation for the ICELL8 protocol.

### Full-length single-cell RNA-seq using ICELL8

The Takara ICELL8 5,184 nano-well chip was used with the full-length SMART-Seq ICELL8 Reagent Kit. Nuclei suspensions were fluorescent-labelled with Hoechst 33,342 for 15 min prior to their dispensing into the Takara ICELL8 5,184 nano-well chips. Cell Select Software (Takara Bio) was used to visualize and select wells containing single nuclei. Nine 5,184 nano-wells chips were used for all samples and 11,084 nuclei were processed for data analysis. Specifically, after quality control, 3,416 nuclei were used for primary PDAC, 3,037 for PDX, and 4,631 for organoids respectively. cDNA synthesis and library preparation were done according to description in previous study ^15^. Libraries were sequenced on the HiSeq 4000 (Illumina) to obtain on average ∼ 0.3 Mio reads per nuclei (SE; 50 bp).

### Bulk-RNA-Seq from primary PDAC samples

Total RNA was extracted from FFPE tumor patient samples using the ReliaPrep™ FFPE Total RNA Miniprep System (Promega). RNA Integrity was determined using the Fragment Analyzer. Because of low RNA integrity (sizing from 50 to 140 bp), we performed a modified TruSeq Stranded Total RNA Library Prep Human/Mouse/Rat (Cat. N°20020596) starting with 200 ng of total RNA. The modifications include a) ignoring fragmentation step, b) ligation optimization by adjusting adapters concentration during library preparation, c) increasing PCR cycles (15 in total) and eliminating primer dimers prior to sequencing (Agencourt AMPure XP magnetic beads, Beckman Coulter). Primary PDAC were sequenced on the NovaSeq6000; S4 flow cell PE 300 cycles generating a data set of 50 to 400 Mio reads per sample.

### Bulk-RNA-Seq from organoids and PDXs

RNA libraries were prepared starting with 300 ng of total RNA using a non-stranded mRNA Seq (TruSeq RNA Library Preparation Cat. RS-122–2001) from Illumina according to the manufacturer’s recommendations. Libraries were sequenced on the Illumina HiSeq 4000 (SE; 1 × 50 bp; 30–35 Mio reads/sample).

### Whole Genome Sequencing

WGS data from two primary PDAC samples TM56 and TM27 were sequenced at ∼40× coverage on the Illumina NovaSeq 6000 sequencer following the protocol provided by the supplier. Libraries were performed using the PCR Free DNA library preparation from Illumina Cat. N°: 20041794). Alignment, variant calling, and benchmarking were performed using Illumina DRAGEN Germline pipeline 4.2.4.

### Pre-processing of single-nuclei RNA-seq data

Raw sequencing files were processed as described in ^12^. Briefly, Cogent NGS analysis pipeline (CogentAP) from Takara Bio (v1.0) was applied for de-multiplexing and creating the gene expression matrices from each FASTQ file. Reads were aligned against the human genome GRCh38 v107 (https://www.ensembl.org/Homo_sapiens/Info/Index). Raw matrices underwent quality-control (QC) filtering for cells and genes as outlined in ^15^ also including intron regions of genes. Gene matrices for cells that passed the quality control were used as input for the SingleCellExperiment R package (v3.0)^49^ to generate SingleCellExperiment objects for the subsequent downstream analysis.

### Identification of disease subtypes and cell types

To identify the tumor subtypes and cell types of which the single cells are composed, two different methods were employed. First, we used the marker-based method AUCell ^30^ to identify the PDAC subtypes classical and basal-like tumor. This analysis was based on established marker genes ^50^. AUCell ranks the genes in each cell by decreasing expression value, and marks cells according to their most expressed marker genes. Secondly, we performed cell type annotation for more refined subpopulations to address the heterogeneity of the tumor and its matched models. We utilized the reference data set as provided by the SeuratData R package (panc8.SeuratData) ^51^ and the prediction function as implemented in the R package SingleR ^52^. Downstream analyses performing the UMAP algorithm were done as implemented in CogentDS (v1.0) for dimensionality reduction and data visualization. To determine which genes were differentially expressed between tumor subtypes and cell types in a particular patient, Wilcoxon Rank Sum and Signed Rank Test was used, together with *p* values adjustment with Benjamini–Hochberg method.

### Pseudo-variant calling in snRNA and bulk RNA-seq data

Bam files resulting from CogentDS were used as input for the pseudo bulk variant calling using GATK best practices pipeline for RNAseq variant calling ^53^. Consistent with the recommendations of GATK, duplicates were removed with Picard MarkDuplicates, and read groups were added with Picard AddOrReplaceReadGroups. Subsequently, Cigar reads were split into exon segment and hard-clip any sequences overhanging into the intronic regions with SplitNCigarReads. Variant calling was performed by HaplotypeCaller and all variants were then hard filtered by the following criteria: FS>30 and QD<2. Patient specific vcf files were intersected or merged using bcftools-isec to collect shared or unique mutations from the samples. Resulting variants were converted into maf (mutation annotation file) using Ensembl VEP ^54^ annotation tool for the identification of the intronic, intergenic, and splice junction mutations (Fig.3-A). Finally, mutations were grouped into three categories: SS, Proximal, and Deep by using Ensembl GTF file version 107. Total number of SS and Prox mutations were subtracted from total number of mutations in maf files to calculate the number of deep intronic variants. Negative values indicate that the deep intronic mutation affects multiple transcripts of the gene. Pre-Ranked enrichment analysis was performed by using mutation lists from three locational groups. Mutation numbers from SS and Proximal regions were normalized by using total intron number of each gene. Deep mutation numbers were normalized by per unit length (kb). Normalized values are ranked and used in enrichment analysis by using R package gProfileR^55^. Overall survival was calculated using Kaplan-Meier analysis based on TCGA-PAAD cohort. The result is shown as Kaplan-Meier plot with p value from log rank test generated by the cBioPortal^56^.

### Integration of the SpliceAI and Pangolin Scoring

To perform scoring on the discovered variant list, hard filtered and intersected VCF files were used. Scores of each variant aggregated on their genes and maximum scores from both algorithms have been taken. Genes that scored with higher than 0.1 one of the tools have been selected as possibly mis-splicing related variants as it stated in their reference manual. SpiceAI v1.3 and Pangolin have been used in our analysis with default parameters with GRCh38 reference genome.

### Cell specific variant calling

To perform single cell specific variant calling, bam files were separated into chip- and sample-specific bam files. Separated bam files were used for the variant calling pipeline, which was applied to individual bam file as before, without the base quality recalibration step (BQSR) to be able to get more variants possible per cell. Quality filtering was applied with the same quality filters as mentioned in pseudo-bulk RNA variant calling in the methods section. VCF files for each barcode were used to collect barcode-specific mutations and visualize as a mutation distribution plot using CogentDS UMAP functions.

### Identification of intron retention events from RNA-Seq data

Intron retention analysis was performed using IRFinder 1.3.0 ^57^. IRFinder calculates IR-ratios to measure IR level reflecting the proportion of intron retaining transcripts. To compare PDAC samples to normal tissue, we downloaded fastq files from nine PDAC and nine healthy human pancreas samples from NBCI GEO database (accession ID: GSE211398). These were pre-processed using the same methods as described in Bulk-RNA-Seq from primary PDAC. Tumor samples from GSE211398 dataset were then analyzed using the IRFinder pipeline to identify IR events. Intronic depth for all samples normalized by CPM due to the library size variations between the public data and CRU samples. Due to the different number of samples between the two dataset, weighted means of IR scores were calculated to balance the variable sample sizes by using 0.1 threshold among 80% of all samples. Normalized intron depths were also used as further filtering to reduce the number of identified possible false negative discoveries.

## Data availability

All sequencing data has been deposited in the Gene Expression Omnibus, Bulk RNA-Sequencing GSE228844 and snRNA-Sequencing GSE229007. Any other relevant data is available from the authors upon reasonable request.

## Code availability

Scripts used to analyze the data and generate the figures and tables in this paper are available on GitHub (https://github.com/UKHG-NIG/DeepVarSplice).

## Supplemental Information

The following supporting information accompanies this manuscript.

**Supplemental Fig. 1:**
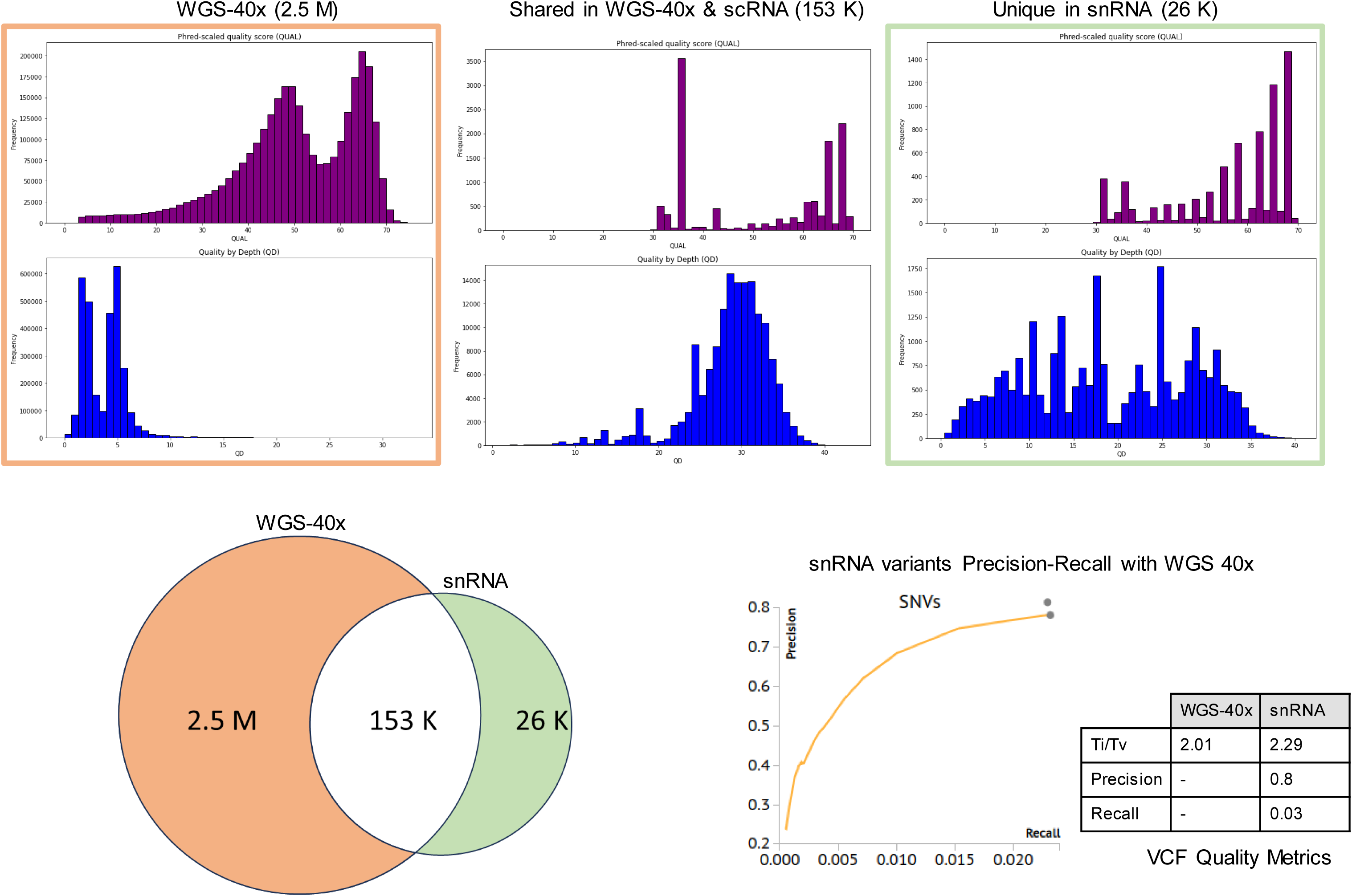
Variant calling quality benchmark obtained by the DeepVarSplice and the gold-standard variant calling (WGS) methods for primary PDACs.

**Supplemental Fig. 2:**
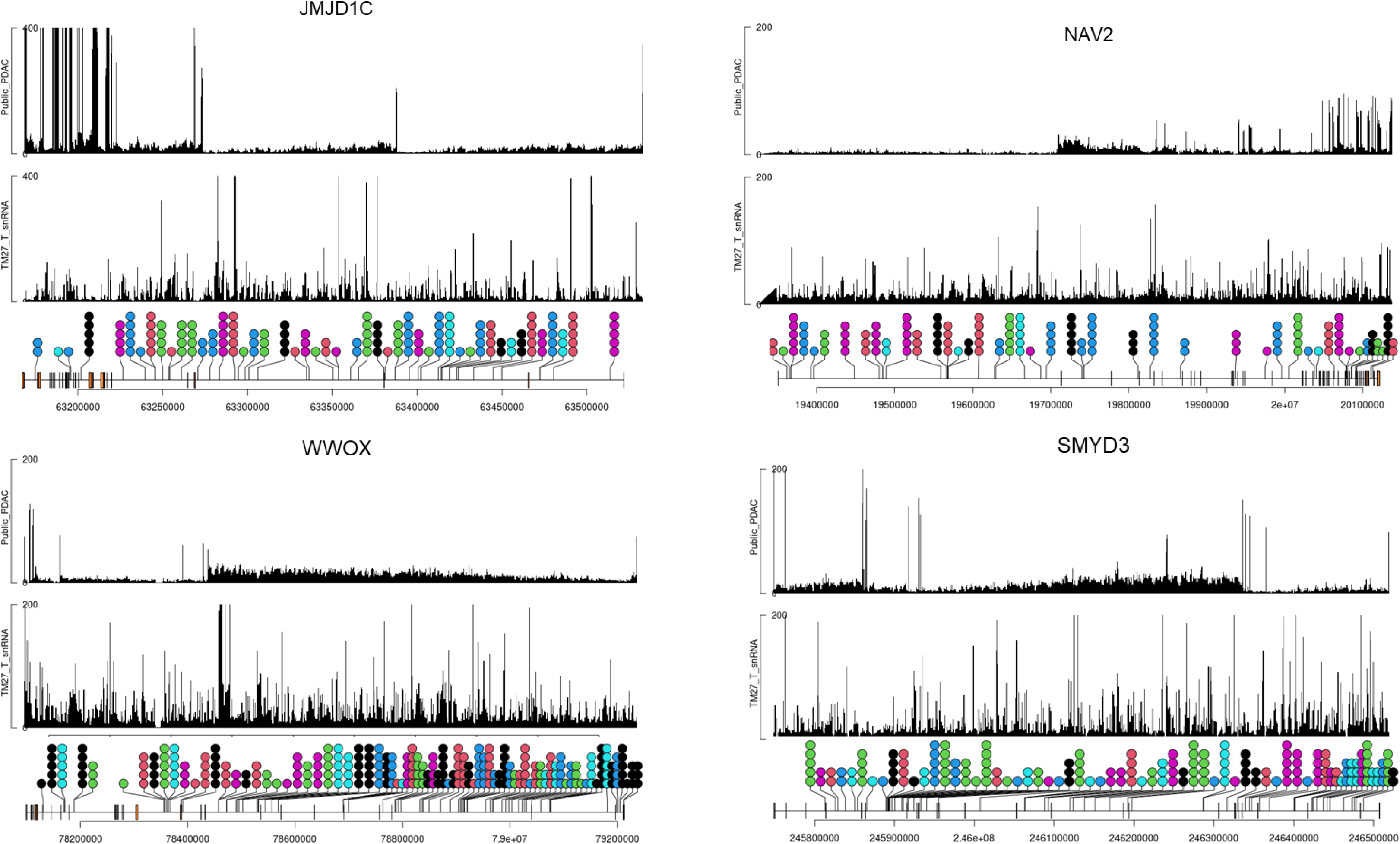
Read coverage of snRNA-seq and bulk RNA-seq data visualizing the intronic variant’s location for *NAV2, JMJCD1, WWOX* and *SMYD3* assessed using pseudo-bulk snRNA-seq data.

**Supplemental Table 1:** Global intronic variants detected from the pseudo-bulk snRNA data.

**Supplemental Table 2:** Number of mutations based on locational classification (Deep-SS-Prox) with normalized numbers by intron length and number.

**Supplemental Table 3:** List of DEGs from classical and basal-like cells obtained from snRNA-seq data of primary PDAC tumors (n=3) and their corresponding PDXs (n=3) and organoids (n=3).

**Supplemental Table 4:** Genes with shared intronic mutations and IR events based on median CPM-normalized reads depth and median IR event occurrence.

## References

1. Shiraishi, Y. et al. Systematic identification of intron retention associated variants from massive publicly available transcriptome sequencing data. Nature Communications 2022 13:1 13, 1–13 (2022).

2. Zhang, P. et al. Genome-wide detection of human variants that disrupt intronic branchpoints. Proc Natl Acad Sci U S A 119, e2211194119–e2211194119 (2022).

3. Park, E., Pan, Z., Zhang, Z., Lin, L. & Xing, Y. The Expanding Landscape of Alternative Splicing Variation in Human Populations. Am J Hum Genet 102, 11–26 (2018).

4. Le Hir, H., Nott, A. & Moore, M. J. How introns influence and enhance eukaryotic gene expression. Trends Biochem Sci 28, 215–220 (2003).

5. Jung, H., Lee, K. S. & Choi, J. K. Comprehensive characterisation of intronic mis-splicing mutations in human cancers. Oncogene 2021 40:7 40, 1347–1361 (2021).

6. Daguenet, E., Dujardin, G. & Valcárcel, J. The pathogenicity of splicing defects: mechanistic insights into pre-mRNA processing inform novel therapeutic approaches. EMBO Rep 16, 1640 (2015).

7. Scotti, M. M. & Swanson, M. S. RNA mis-splicing in disease. Nat Rev Genet 17, 19–32 (2016).

8. Qian, X. et al. Identification of Deep-Intronic Splice Mutations in a Large Cohort of Patients with Inherited Retinal Diseases. Front Genet 12, 276 (2021).

9. Sahin, I., George, A. & Seyhan, A. A. Therapeutic Targeting of Alternative RNA Splicing in Gastrointestinal Malignancies and Other Cancers. Int J Mol Sci 22, (2021).

10. Raguraman, P., Balachandran, A. A., Chen, S., Diermeier, S. D. & Veedu, R. N. Antisense Oligonucleotide-Mediated Splice Switching: Potential Therapeutic Approach for Cancer Mitigation. Cancers (Basel) 13, (2021).

11. Peng, Q. et al. Impacts and mechanisms of alternative mRNA splicing in cancer metabolism, immune response, and therapeutics. Molecular Therapy 30, 1018–1035 (2022).

12. Bonnal, S. C., López-Oreja, I. & Valcárcel, J. Roles and mechanisms of alternative splicing in cancer - implications for care. Nat Rev Clin Oncol 17, 457–474 (2020).

13. Tress, M. L., Abascal, F. & Valencia, A. Alternative Splicing May Not Be the Key to Proteome Complexity. Trends Biochem Sci 42, 98–110 (2017).

14. Wang, X., He, Y., Zhang, Q., Ren, X. & Zhang, Z. Direct Comparative Analyses of 10X Genomics Chromium and Smart-seq2. Genomics Proteomics Bioinformatics 19, 253–266 (2021).

15. Shomroni, O. et al. A novel single-cell RNA-sequencing approach and its applicability connecting genotype to phenotype in ageing disease. Sci Rep 12, 4091–4091 (2022).

16. Kuo, H. C. et al. A Comparative Proteomic Analysis of Erinacine A’s Inhibition of Gastric Cancer Cell Viability and Invasiveness. Cellular Physiology and Biochemistry 43, 195–208 (2017).

17. Huang, T. et al. Csmd1 mutations are associated with increased mutational burden, favorable prognosis, and anti-tumor immunity in gastric cancer. Genes (Basel) 12, (2021).

18. Chen, H. et al. Association of LRP1B mutation with tumor mutation burden and outcomes in melanoma and non-small cell lung cancer patients treated with immune check-point blockades. Front Immunol 10, (2019).

19. Datta, N., Chakraborty, S., Basu, M. & Ghosh, M. K. Tumor Suppressors Having Oncogenic Functions: The Double Agents. Cells 10, 1–26 (2021).

20. Del Mare, S., Salah, Z. & Aqeilan, R. I. WWOX: its genomics, partners, and functions. J Cell Biochem 108, 737–745 (2009).

21. Salah, Z., Aqeilan, R. & Huebner, K. WWOX gene and gene product: tumor suppression through specific protein interactions. Future Oncol 6, 249 (2010).

22. Iliopoulos, D. et al. Roles of FHIT and WWOX fragile genes in cancer. Cancer Lett 232, 27–36 (2006).

23. Singh, S.K., Kumar, S., Viswakarma, N. et al. MAP4K4 promotes pancreatic tumorigenesis via phosphorylation and activation of mixed lineage kinase 3. Oncogene 40, 6153–6165 (2021).

24 Sun, S., Zhang, Z., Fregoso, O. & Krainer, A. R. Mechanisms of activation and repression by the alternative splicing factors RBFOX1/2. RNA 18, 274–283 (2012).

25. Lin, F. et al. Identification of inflammatory response and alternative splicing in acute kidney injury and experimental verification of the involvement of RNAbinding protein RBFOX1 in this disease. Int J Mol Med 49, (2022).

26. Jbara, A. et al. RBFOX2 modulates a metastatic signature of alternative splicing in pancreatic cancer. Nature (2023) doi:10.1038/S41586-023-05820-3.

27. Cowling, V. H. Regulation of mRNA cap methylation. Biochem J 425, 295–302 (2009).

28. Jaganathan K, et al. Predicting Splicing from Primary Sequence with Deep Learning. Cell 176 (3):535–548 (2019).

29. Zeng, T., Li, Y.I. Predicting RNA splicing from DNA sequence using Pangolin. Genome Biol 23, 103 (2022).

30. Aibar, S. et al. SCENIC: single-cell regulatory network inference and clustering. Nature Methods 2017 14:11 14, 1083–1086 (2017).

31. Xiong, D. D. et al. The clinical value of lncRNA NEAT1 in digestive system malignancies: A comprehensive investigation based on 57 microarray and RNA-seq datasets. Oncotarget 8, 17665 (2017).

32. Duxbury, M. S. et al. CEACAM6 is a novel biomarker in pancreatic adenocarcinoma and PanIN lesions. Ann Surg 241, 491–496 (2005).

33. Striefler, J. K. et al. Mucin-1 Protein Is a Prognostic Marker for Pancreatic Ductal Adenocarcinoma: Results From the CONKO-001 Study. Front Oncol 11, 1 (2021).

34. Kumar, S. & Mishra, S. MALAT1 as master regulator of biomarkers predictive of pan-cancer multi-drug resistance in the context of recalcitrant NRAS signaling pathway identified using systems-oriented approach. Sci Rep 12, (2022).

35. Wong, J. J. L., Au, A. Y. M., Ritchie, W. & Rasko, J. E. J. Intron retention in mRNA: No longer nonsense: Known and putative roles of intron retention in normal and disease biology. Bioessays 38, 41–49 (2016).

36. Jung, H. et al. Intron retention is a widespread mechanism of tumor-suppressor inactivation. Nature Genetics 2015 47:11 47, 1242–1248 (2015).

37. Ge, Y. & Porse, B. T. The functional consequences of intron retention: alternative splicing coupled to NMD as a regulator of gene expression. Bioessays 36, 236–243 (2014).

38. Pandit, S. et al. Genome-wide analysis reveals SR protein cooperation and competition in regulated splicing. Mol Cell 50, 223–235 (2013).

39. Davis, R. L., Homer, V. M., George, P. M. & Brennan, S. O. A deep intronic mutation in FGB creates a consensus exonic splicing enhancer motif that results in afibrinogenemia caused by aberrant mRNA splicing, which can be corrected in vitro with antisense oligonucleotide treatment. Hum Mutat 30, 221–227 (2009).

40. Homolova, K. et al. The deep intronic c.903+469T>C mutation in the MTRR gene creates an SF2/ASF binding exonic splicing enhancer, which leads to pseudoexon activation and causes the cblE type of homocystinuria. Hum Mutat 31, 437–444 (2010).

41. Vo, T. et al. HNRNPH1 destabilizes the G-quadruplex structures formed by G-rich RNA sequences that regulate the alternative splicing of an oncogenic fusion transcript. Nucleic Acids Res 50, 6474–6496 (2022).

42. Murray, J. I., Voelker, R. B., Henscheid, K. L., Warf, M. B. & Berglund, J. A. Identification of motifs that function in the splicing of non-canonical introns. Genome Biol 9, 1–17 (2008).

43. Brouard, JS., Schenkel, F., Marete, A. et al. The GATK joint genotyping workflow is appropriate for calling variants in RNA-seq experiments. J Animal Sci Biotechnol 10, 44 (2019).

44. Grellscheid, S.-N. & Smith, C. W. J. An Apparent Pseudo-Exon Acts both as an Alternative Exon That Leads to Nonsense-Mediated Decay and as a Zero-Length Exon. Mol Cell Biol 26, 2237 (2006).

45. Han, K., Yeo, G., An, P., Burge, C. B. & Grabowski, P. J. A Combinatorial Code for Splicing Silencing: UAGG and GGGG Motifs. PLoS Biol 3, e158 (2005).

46. Singh, R. RNA–Protein Interactions That Regulate Pre-mRNA Splicing. Gene Expr 10, 79 (2002).

47. Rahman, M. A., Lin, K. T., Bradley, R. K., Abdel-Wahab, O. & Krainer, A. R. Recurrent SRSF2 mutations in MDS affect both splicing and NMD. Genes Dev 34, 413–427 (2020).

48. Jankowsky, E. & Harris, M. E. Specificity and nonspecificity in RNA–protein interactions. Nature Reviews Molecular Cell Biology 2015 16:9 16, 533–544 (2015).

49. Amezquita, R. A. et al. Orchestrating single-cell analysis with Bioconductor. Nature Methods 2019 17:2 17, 137–145 (2019).

50. Chan-Seng-Yue, M. et al. Transcription phenotypes of pancreatic cancer are driven by genomic events during tumor evolution. Nature Genetics 2020 52:2 52, 231–240 (2020).

51. Segerstolpe, Å. et al. Single-Cell Transcriptome Profiling of Human Pancreatic Islets in Health and Type 2 Diabetes. Cell Metab 24, 593–607 (2016).

52. Aran, D. et al. Reference-based analysis of lung single-cell sequencing reveals a transitional profibrotic macrophage. Nature Immunology 2019 20:2 20, 163–172 (2019).

53. Van der Auwera, G. A. et al. From FastQ Data to High-Confidence Variant Calls: The Genome Analysis Toolkit Best Practices Pipeline. Curr Protoc Bioinformatics 43, 11.10.1–11.10.33 (2013).

54. McLaren, W. et al. The Ensembl Variant Effect Predictor. Genome Biol 17, 1–14 (2016).

55. Peterson, H., Kolberg, L., Raudvere, U., Kuzmin, I. & Vilo, J. gprofiler2 -- an R package for gene list functional enrichment analysis and namespace conversion toolset g:Profiler. F1000Research 2020 9:709 9, 709 (2020).

56. Gao, J. et al. Integrative analysis of complex cancer genomics and clinical profiles using the cBioPortal. Sci Signal 6, (2013).

57. Middleton, R. et al. IRFinder: Assessing the impact of intron retention on mammalian gene expression. Genome Biol 18, 1–11 (2017).

